# Visual system structural and functional connections during face viewing in body dysmorphic disorder

**DOI:** 10.1101/2024.07.12.603273

**Authors:** Wan-wa Wong, Hayden Peel, Ryan Cabeen, Joel P. Diaz-Fong, Jamie D. Feusner

## Abstract

**Background:** Individuals with body dysmorphic disorder (BDD) perceive distortions in their appearance, which could be due to imbalances in global and local visual processing. The vertical occipital fasciculus connects dorsal and ventral visual stream regions, integrating global and local information, yet the role of this structural connection in BDD has not been explored. Here, we investigated the vertical occipital fasciculus’s white matter microstructure in those with BDD and healthy controls and tested associations with psychometric measures and effective connectivity while viewing their face during fMRI.

**Methods:** We analyzed diffusion MRI and fMRI data in 17 unmedicated adults with BDD and 21 healthy controls. For diffusion MRI, bundle-specific analysis was performed, enabling quantitative estimation of neurite density and orientation dispersion of the vertical occipital fasciculus. For task fMRI, participants naturalistically viewed photos of their own face, from which we computed effective connectivity from dorsal to ventral visual regions.

**Results:** In BDD, neurite density was negatively correlated with appearance dissatisfaction and negatively correlated with effective connectivity. Further, those with weaker effective connectivity while viewing their face had worse BDD symptoms and worse insight. In controls, no significant relationships were found between any of the measures. There were no significant group differences in neurite density or orientation dispersion.

**Conclusion:** Those with BDD with worse appearance dissatisfaction have a lower fraction of tissue having axons or dendrites along the vertical occipital fasciculus bundle, possibly reflecting impacting the degree of integration of global and local visual information between the dorsal and ventral visual streams. These results provide early insights into how the vertical occipital fasciculus’s microstructure relates to the subjective experience of one’s appearance, as well as the possibility of distinct functional-structural relationships in BDD.

## 1 Introduction

Individuals with body dysmorphic disorder (BDD) are preoccupied with misperceived appearance defects, which they believe render them ugly and deformed. Misperceived appearance features in BDD commonly involve the face and head but can be of any part of the body^[1]^. High lifetime rates of suicide attempts (25%)^[2]^ and hospitalization (50%)^[3]^ underscore its potential severity, and it has a relatively high point prevalence of ∼2% in the general population^[4]^. Yet, BDD is often misdiagnosed, or the diagnosis is missed, and it is still understudied. While some neurobiological models have been put forth to explain the formation and maintenance of symptoms in BDD ^[5, 6]^, a comprehensive understanding of this condition is still needed.

Aberrant visual information processing likely contributes to appearance misperceptions experienced by those with BDD ^[6, 7]^. Previous functional MRI (fMRI) studies, using own-face^[8]^, other-face^[9]^, and house^[10]^ stimuli found reduced activity in BDD compared with controls in the dorsal visual stream (DVS) when viewing filtered images that contained only low spatial frequency (LSF) information. (Spatial frequency refers to the number of brightness changes (i.e., cycles) per 1° of visual angle.) LSF images emphasize global properties/contours of images and are processed predominantly in the magnocellular pathway extending through the DVS, whereas high spatial frequency (HSF) images emphasize details and other high image contrast features and are processed predominantly in the parvocellular pathway extending through the ventral visual stream (VVS)^[11]^.

These findings of reduced DVS activity in BDD when viewing the filtered images have contributed to a model in which the hyper-scrutiny of miniscule appearance details could be mechanistically related to failing to “see” the appearance feature as an integrated whole, which may reflect an imbalance in global and local visual processing. Several lines of evidence support this hypothesis. Psychophysical and neurophysiological evidence has demonstrated that robust phenomena like the face and body inversion effects, which rely more heavily on featural as opposed to global processing, are less pronounced in BDD and those with high body dysmorphic symptoms compared to healthy controls ^[12–17]^. Imaging and electro-cortical studies subsequently demonstrated DVS hypoactivity when viewing LSF images ^[18, 19]^. Further, in one study, VVS hyperactivity in BDD compared to controls when viewing HSF (high-detail) images was found, along with significant associations between the magnitude of VVS activation and the perception of faces as less attractive^[18]^. Functional connectivity studies align with these results. During an others’ face-viewing task, individuals with BDD demonstrated hyperconnectivity in the VVS compared to healthy controls between left anterior occipital face area and right fusiform face area (FFA) for LSF faces^[20]^. In another study, during a body-viewing task, BDD demonstrated hypoconnectivity in the dorsal visual network^[21]^. These various lines of evidence converge and have helped form a theory of functional global/local visual processing imbalances in BDD ^[22]^.

However, the structural connections underlying functional communications between visual regions have been less explored. The vertical occipital fasciculus (VOF) is a fibre bundle that runs principally superior-inferior within the occipital lobe ^[23, 24]^. The VOF may play a role in integrating information between the DVS, involved in the eye position signals, attention and motion perception^[25–27]^, and the VVS, involved in part-based processing and visual categorization (e.g. words, faces and bodies) ^[28, 29]^. This is an important connection, as information from the DVS and VVS converge in inferior temporal regions such that the more rapidly-processed global and configural information, as a function of the DVS, provides a template for contextualization of the more slowly-processed details arriving from the ventral stream [30-32]. Of note, the FFA, involved in recognition of own and others’ faces ^[33]^, is in the VVS [34] in inferior temporal regions that additionally receive DVS input ^[29, 35]^. However, no study has specifically investigated the VOF in BDD. Characterizing the white matter (WM) microstructure of the VOF from diffusion MRI (dMRI) data, relationships with clinical variables, as well as the functional connections between corresponding DVS and VVS regions have not been examined. Exploring these relationships may provide a better understanding of potential neuropathological mechanisms related to dorsal/ventral visual system information transfer that could underlie perceptual distortions and related symptoms in BDD.

Here, we investigated the WM microstructure of the VOF, estimated with neurite orientation dispersion and density imaging (NODDI) indices. NODDI is a diffusion MRI technique for the estimation of key aspects of neuronal tissue for each voxel. NODDI measures include neurite density index (NDI), which quantifies the packing density of axons or dendrites; and orientation dispersion index (ODI), which assesses the orientational coherence of neurites ^[36]^. Such indices of neurites provide more specific markers of brain tissue microstructure than standard indices from diffusion tensor imaging, such as fractional anisotropy (FA), which describes the degree of anisotropy of water molecules, and mean diffusivity (MD), which measures the overall diffusivity in the tissue ^[36]^. We compared these indices (NDI and ODI) along the VOF between groups and examined their associations with psychometric measures and with dynamic effective connectivity (DEC) from the DVS to the VVS during fMRI while participants viewed pictures of their own face. In line with the imbalanced global and local processing model^[22]^, we hypothesized differences in VOF WM microstructure in BDD compared to healthy controls which may signify disrupted integration between regions involved in global and local visual processing. We also hypothesized WM microstructure metrics would be associated with appearance dissatisfaction and BDD symptom severity in BDD and with appearance dissatisfaction in healthy controls. Relationships between WM microstructure metrics of the VOF and DEC between the dorsal and ventral streams; between DEC and clinical variables; and among WM, DEC, and clinical variables; were investigated in an exploratory manner.

## 2 Materials and Methods

### 2.1 Participants

The UCLA Institutional Review Board approved the study. All participants provided informed written consent. Twenty-one unmedicated adults with BDD and 23 healthy controls (CON) aged 18-40 years were recruited from the community and were enrolled. BDD participants met DSM-5 criteria for BDD, with face concerns. BDD participants could have comorbid depressive or anxiety disorders, since they commonly co-occur, but not other comorbid psychiatric or substance use disorders (See Supplementary Material S1 for full inclusion and exclusion criteria).

### 2.2 Clinical assessments

Eligibility was determined through telephone screening followed by a clinical interview with the study physician (JDF). The Mini International Neuropsychiatric Interview (MINI) and BDD Module ^[37, 38]^ were administered to all participants. The Yale-Brown Obsessive-Compulsive Scale Modified for BDD (BDD-YBOCS) ^[39]^, measuring BDD-related preoccupations/repetitive behaviours, was administered to BDD participants. The Body Image States Scale (BISS) ^[40]^, measuring appearance dissatisfaction (lower scores indicating a greater degree of appearance dissatisfaction), was administered to BDD and CON participants. The Brown Assessment of Beliefs Scale (BABS), measuring BDD-related insight, was administered to BDD participants and the Montgomery-Åsberg Depression Rating Scale (MADRS) [41] and the Hamilton Anxiety Scale (HAMA) ^[42]^ were administered to BDD and CON participants (See Supplementary Material S2 for assessment details).

### 2.3 MRI data acquisition and processing

MRI data were acquired on a 3T Siemens Prisma scanner. Diffusion MRI (dMRI) data processing was done using a combination of FSL diffusion toolbox ^[43]^ and quantitative imaging toolkit (QIT) ^[44]^, and functional MRI data processing was done using fMRIPrep^[45]^. See Supplementary Material S3-S5 for details of data acquisition and processing, including quality control and motion correction.

### 2.4 WM microstructure analysis

Reconstruction of white matter tracts was done using tractography to obtain microstructure maps of the VOF in the left and right hemispheres (see Fig. 1). Geometric models of WM connectivity were reconstructed from fibre orientation data estimated from dMRI. A bundle-specific analysis was then performed, enabling the quantitative estimation of NODDI ^[36]^ (NDI and ODI) indices of the whole bundle.

**Figure 1.**
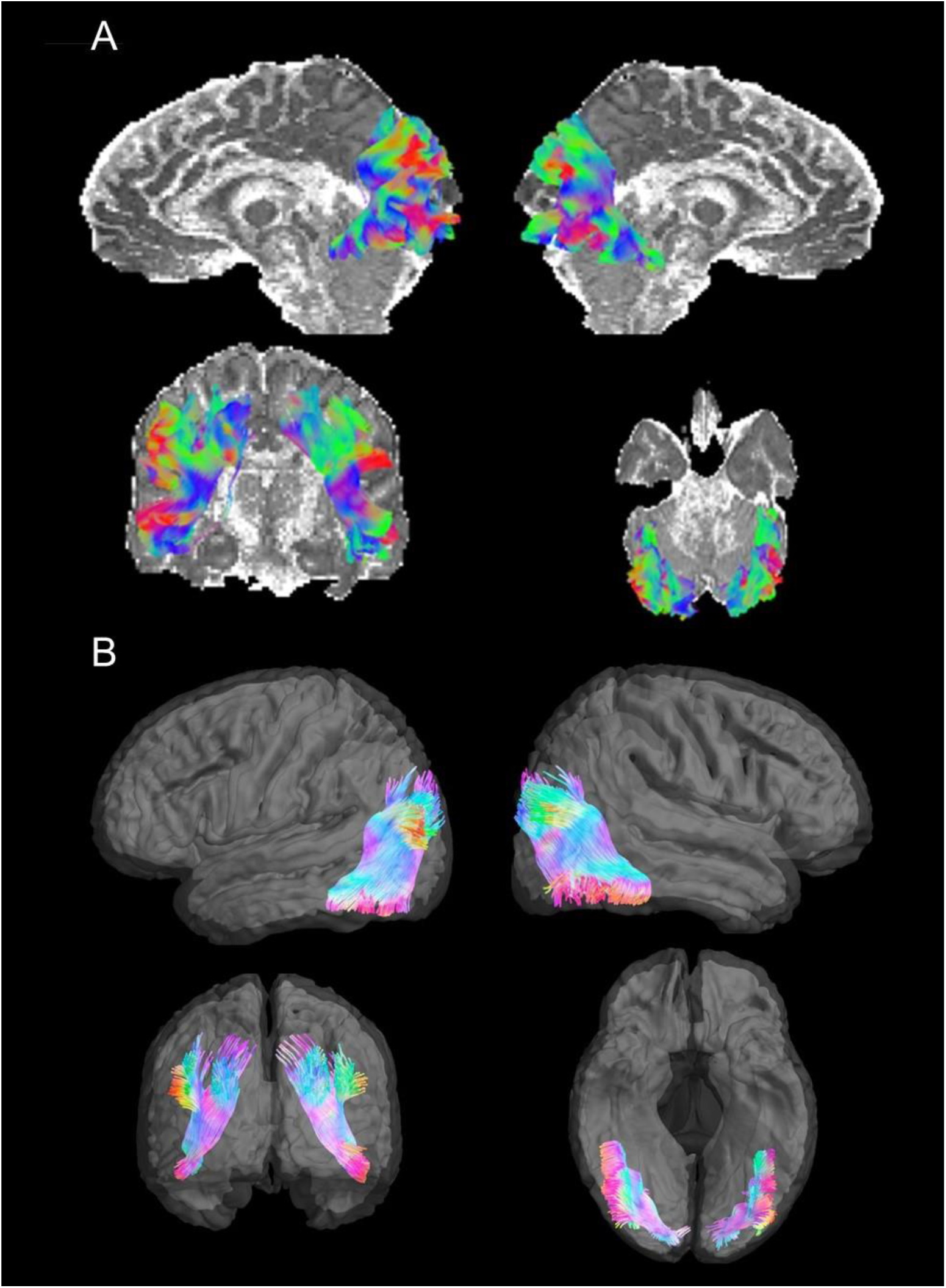
VOF tractography fibre reconstruction in one representative BDD participant (A), and the probabilistic VOF reconstruction from the average template within the QIT software (B)

We averaged the point mean values (i.e. the means among the bundle vertices [points]) of these indices of the bilateral VOF for subsequent analyses.

### 2.5 fMRI task paradigm

The fMRI task paradigm was performed as previously described ^[46, 47]^. We used the first, self-face “natural viewing” task and the control task in the present analysis. There were two sets of stimuli for the natural viewing condition: photos of the participant’s face and scrambled faces as the control task. Participants were instructed to view the (unaltered) photos of their face and scrambled images of their face as they normally do. They were instructed to press a button every time an image disappeared from the screen to ensure vigilance.

### 2.6 Brain functional connectivity analysis

Regions-of-interest (ROIs) in the DVS and VVS were derived from the Neurosynth functional meta-analysis (https://neurosynth.org/) with the search terms “primary visual,” “ventral visual,” “visual stream,” and “dorsal visual” to obtain maps generated with association tests, as previously described^[47]^. Blind-deconvolution^[48]^ was performed on the time series extracted from these ROIs to minimize intra-subject variability in hemodynamic response function (HRF) ^[49]^, and to improve estimation of effective connectivity^[50]^. DEC, a time-varying measure of directional connectivity between pairs of ROIs, was computed at each time point using time-varying Granger causality (GC)^[51]^. DEC was used because of its ability to estimate causal connectivity across time with a precision of each timepoint, which helped us capture connectivity only within task blocks of interest. The deconvolved timeseries were fitted into a dynamic multivariate autoregressive (dMVAR) model for estimating DEC between ROIs, which was solved in a Kalman-filter framework. The dMVAR model coefficients vary as a function of time, whose lengths were identical to the number of timepoints in the time series (see Supplementary Material S6 for additional information). Eight total intra-hemispheric connections (see Fig. 2) were chosen between DVS (inferior parietal lobule [IPL] and superior parietal lobule [SPL]) and VVS (inferior temporal gyrus [ITG] and fusiform gyrus [FG]) ROIs: DVS to VVS *(ipsilateral IPL to FG; ipsilateral IPL to ITG; ipsilateral SPL to FG; ipsilateral SPL to ITG)* in both hemispheres. From these connections, we extracted the timepoints associated with trials involving unaltered faces for subsequent statistical analysis. To limit the number of comparisons because of the sample size and because we did not have specific a priori hypotheses about which hemisphere or connections would exhibit differences in DEC, or associations with clinical variables, we averaged the eight connections.

**Figure 2.**
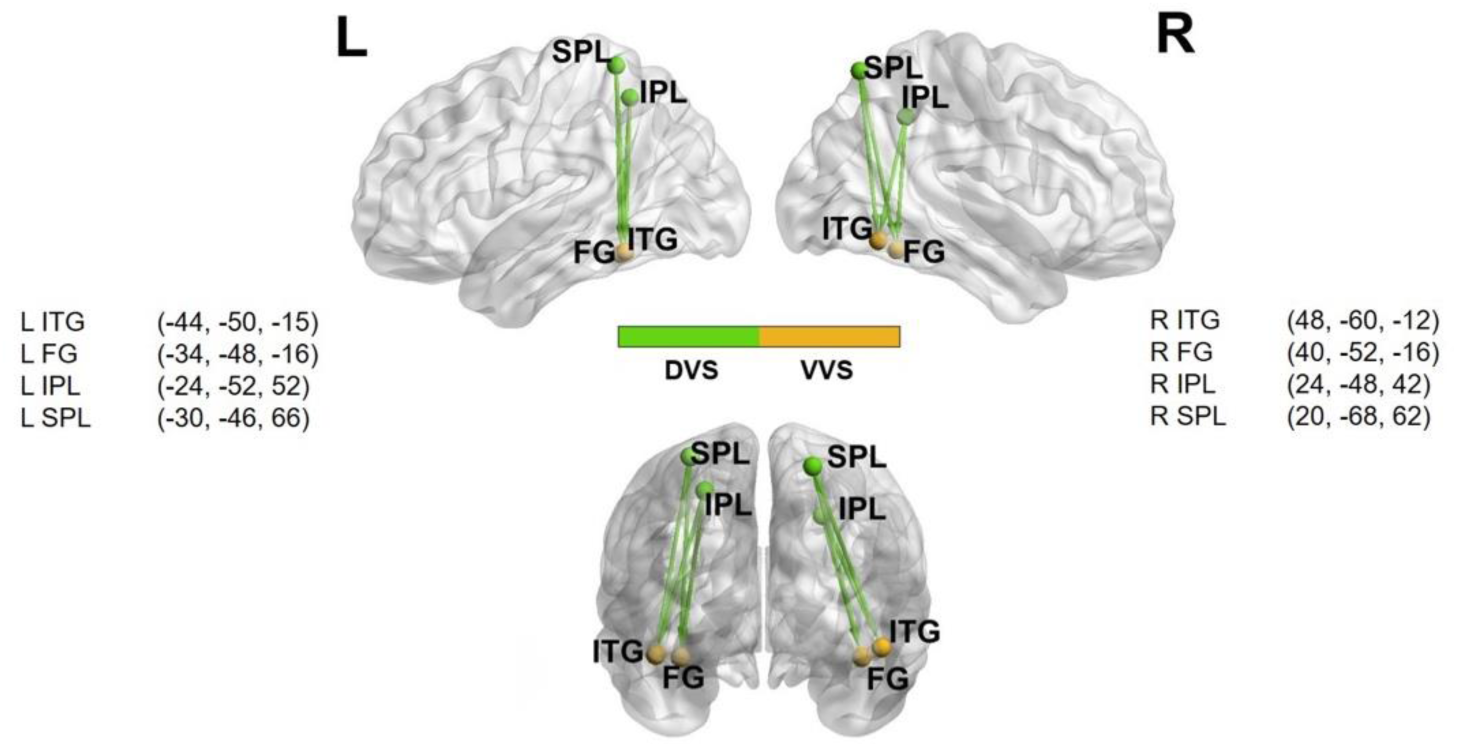
Locations of the eight spherical ROIs used for DEC analyses, overlaid on a brain surface with posterior and lateral views. Four were in the DVS [bilateral superior parietal lobule (SPL) and bilateral inferior parietal lobule (IPL)] and four were in the VVS [bilateral inferior temporal gyrus (ITG) and bilateral fusiform gyrus (FG)]. Note the directional connections between DVS and VVS ROIs. All spheres had a radius of 5 mm and the center-of-mass coordinates obtained from the clusters in x, y, and z in the MNI space. This panel was prepared using BrainNet Viewer ^[52]^.

### 2.7 Statistical analyses

Independent-samples *t*-tests were used to test for significant group differences. Pearson correlations were used to determine associations between the hypothesized symptom severity measures (BDD-YBOCS and BISS) and VOF WM microstructure and, in exploratory analyses, DEC and BABS. We used alpha = .05 for a statistical significance threshold. See Supplementary Tables 1-3 for assumption testing. Because we tested two WM microstructural elements without specific hypotheses for each separately, to reduce the chance of false discoveries, we used Bonferroni correction for the group comparisons and the Benjamini-Hochberg False Discovery Rate (FDR) method for correlation analyses.

We additionally performed *post hoc* moderation analyses using Andrew Hayes’ Process macro (version 4.2)[53]. Moderation occurs when the relationship between two variables depends on a third variable (i.e., the moderator). These analyses were conducted to explore whether DEC moderates -- i.e., affects the direction/strength of the relationship between -- VOF WM microstructure and BISS. We conducted statistical tests in SPSS and R.

## 3 Results

### 3.1 Sample characteristics

Twenty-one BDD participants and 23 CON were eligible who completed the entire protocol (Table 1). Of these, we excluded three BDD and two CON participants’ data due to excessive motion artifacts, and one BDD participant’s data due to fMRIPrep errors. Therefore, fMRI data from 17 BDD and 21 CON were included in subsequent analyses.

**Table 1.**
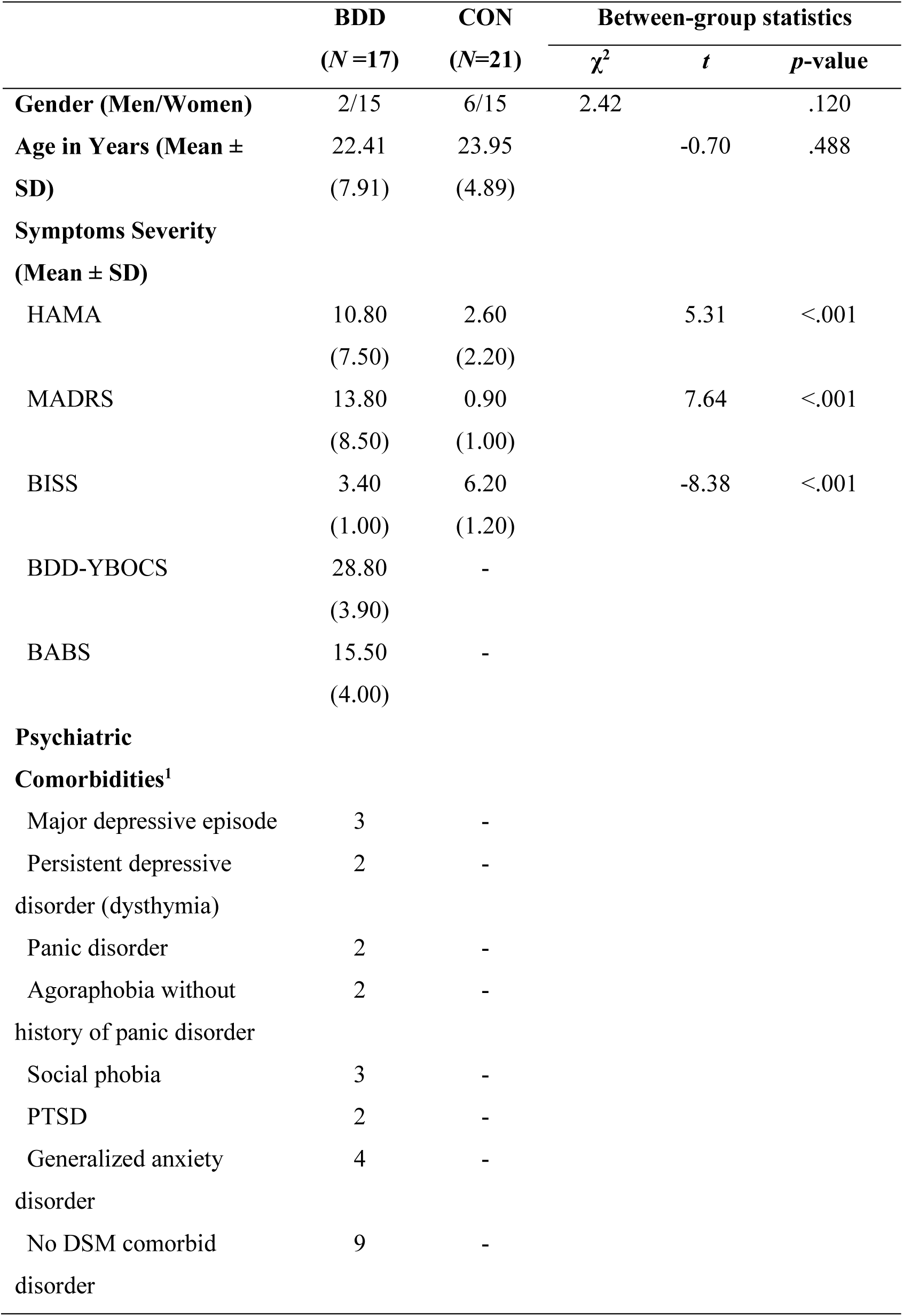
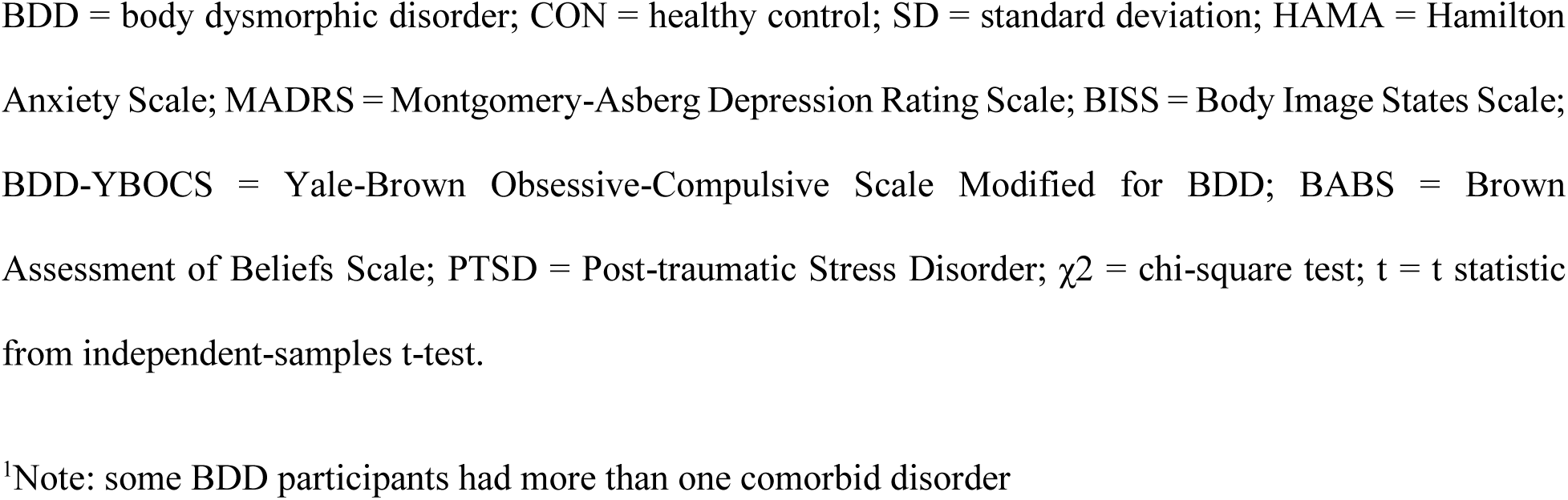
Demographics and Psychometrics.

|  | **BDD**<br>(*N* =17) | **CON**<br>(*N*=21) | **Between-group statistics** |  |  |
| --- | --- | --- | --- | --- | --- |
| | | | $\chi^2$ | *t* | *p*-value |
| **Gender (Men/Women)** | 2/15 | 6/15 | 2.42 |  | .120 |
| **Age in Years (Mean ± SD)** | 22.41<br>(7.91) | 23.95<br>(4.89) |  | -0.70 | .488 |
| **Symptoms Severity (Mean ± SD)** |  |  |  |  |  |
| HAMA | 10.80<br>(7.50) | 2.60<br>(2.20) |  | 5.31 | <.001 |
| MADRS | 13.80<br>(8.50) | 0.90<br>(1.00) |  | 7.64 | <.001 |
| BISS | 3.40<br>(1.00) | 6.20<br>(1.20) |  | -8.38 | <.001 |
| BDD-YBOCS | 28.80<br>(3.90) | - |  |  |  |
| BABS | 15.50<br>(4.00) | - |  |  |  |
| **Psychiatric Comorbidities<sup>1</sup>** |  |  |  |  |  |
| Major depressive episode | 3 | - |  |  |  |
| Persistent depressive disorder (dysthymia) | 2 | - |  |  |  |
| Panic disorder | 2 | - |  |  |  |
| Agoraphobia without history of panic disorder | 2 | - |  |  |  |
| Social phobia | 3 | - |  |  |  |
| PTSD | 2 | - |  |  |  |
| Generalized anxiety disorder | 4 | - |  |  |  |
| No DSM comorbid disorder | 9 | - |  |  |  |

BDD = body dysmorphic disorder; CON = healthy control; SD = standard deviation; HAMA = Hamilton Anxiety Scale; MADRS = Montgomery-Asberg Depression Rating Scale; BISS = Body Image States Scale; BDD-YBOCS = Yale-Brown Obsessive-Compulsive Scale Modified for BDD; BABS = Brown Assessment of Beliefs Scale; PTSD = Post-traumatic Stress Disorder; $\chi^2$ = chi-square test; t = t statistic from independent-samples t-test.
<sup>1</sup>Note: some BDD participants had more than one comorbid disorder

### 3.2 Group differences in WM microstructure and dynamic connectivity

At an uncorrected significance level, the BDD group demonstrated lower mean ODI – smaller angular variation and of neurite structures – compared to controls (*p*=.043); however, this did not survive Bonferroni corrections for multiple comparisons (.05/2 = .025). No significant group difference was detected in DEC (See Table 2).

**Table 2.** Group comparisons in WM microstructure and dynamic connectivity.

|  | BDD | CON | Between-group statistics |  |  |
| --- | --- | --- | --- | --- | --- |
| | ($M \pm SD$) | ($M \pm SD$) | $t$ | $p$-value | Cohen's D [95% CI] |
| **NDI** | 0.608 ±<br>0.021 | 0.602 ±<br>0.027 | 0.77 | .447 | 0.248 [-0.40, 0.89] |
| **ODI** | 0.245 ±<br>0.009 | 0.252 ±<br>0.011 | -2.09 | .043 | -0.683 [-1.34, -0.02] |
| **DEC** | 0.028 ±<br>0.070 | 0.017 ±<br>0.059 | 0.53 | .603 | 0.172 [-0.47, 0.81] |

BDD = body dysmorphic disorder; CON = control; NDI = neurite density index; ODI = orientation dispersion index; DEC = dynamic effective connectivity; $t$ = independent-samples $t$ -test.

### 3.3 Relationships of clinical symptoms with WM microstructure and dynamic connectivity

#### 3.3.1 Testing hypotheses

In BDD, NDI of the VOF was significantly, positively correlated with BISS (*r* = .68, 95% CI = [.30, .88], *p* = .003, FDR adj. *p* = .012), and ODI of the VOF was positively correlated with BDD-YBOCS, although this did not survive corrections for multiple comparisons (*r* = .48, *p* = .050, FDR adj. *p* = .100). Other relationships between NDI and BDD-YBOCS (*r* = .25, *p* = .338, FDR adj. *p* = .357), ODI and BISS (*r* = −.24, *p* = .357, FDR adj. *p* = .357) were non-significant. In CON, no significant correlations were found between NDI or ODI and BISS scores (*r* = .31, *p* = .172, and *r* = .26, *p* = .255, respectively).

#### 3.3.2 Exploratory analyses

In BDD, NDI of the VOF was significantly, negatively correlated with DEC (*r*= −.61, 95% CI = [−.84, −.18], *p* = .009, FDR adj. *p* = .030), and positively correlated with BABS (*r* = .51, *p* = .035, FDR adj. *p* = .053) although the latter did not survive correction for multiple comparisons. No significant correlation was detected between ODI and DEC (*r* = −.18, *p* = .492, FDR adj. *p* = .492). In CON, there was a trend for ODI being negatively correlated with DEC (*r* = −0.41, *p* = 0.068, FDR adj. *p* = 204), but it did not survive corrections for multiple comparisons. There was also no significant relationship between NDI and DEC (*r* = .01, *p* = .984, FDR adj. *p* = .984).

In BDD, there was a significant negative correlation between BDD-YBOCS and DEC (*r* = −.58, 95% CI [−.83, −.13], *p* = .015, FDR adj. *p* = .030), and between BABS and DEC (*r* = −.59, 95% CI [−.84, −.16], *p* =.012, FDR adj. *p* = .030); those with weaker connectivity have worse BDD symptoms and poorer insight. In BDD and healthy controls, there were no significant correlations between DEC and BISS (BDD: *r* = −.38, *p* = .137, FDR adj. *p* = .210; CON: *r* = −0.26, *p* = .261, FDR adj. *p* = .392). See Fig. 3 for visualization of all significant correlations.

**Figure 3.**
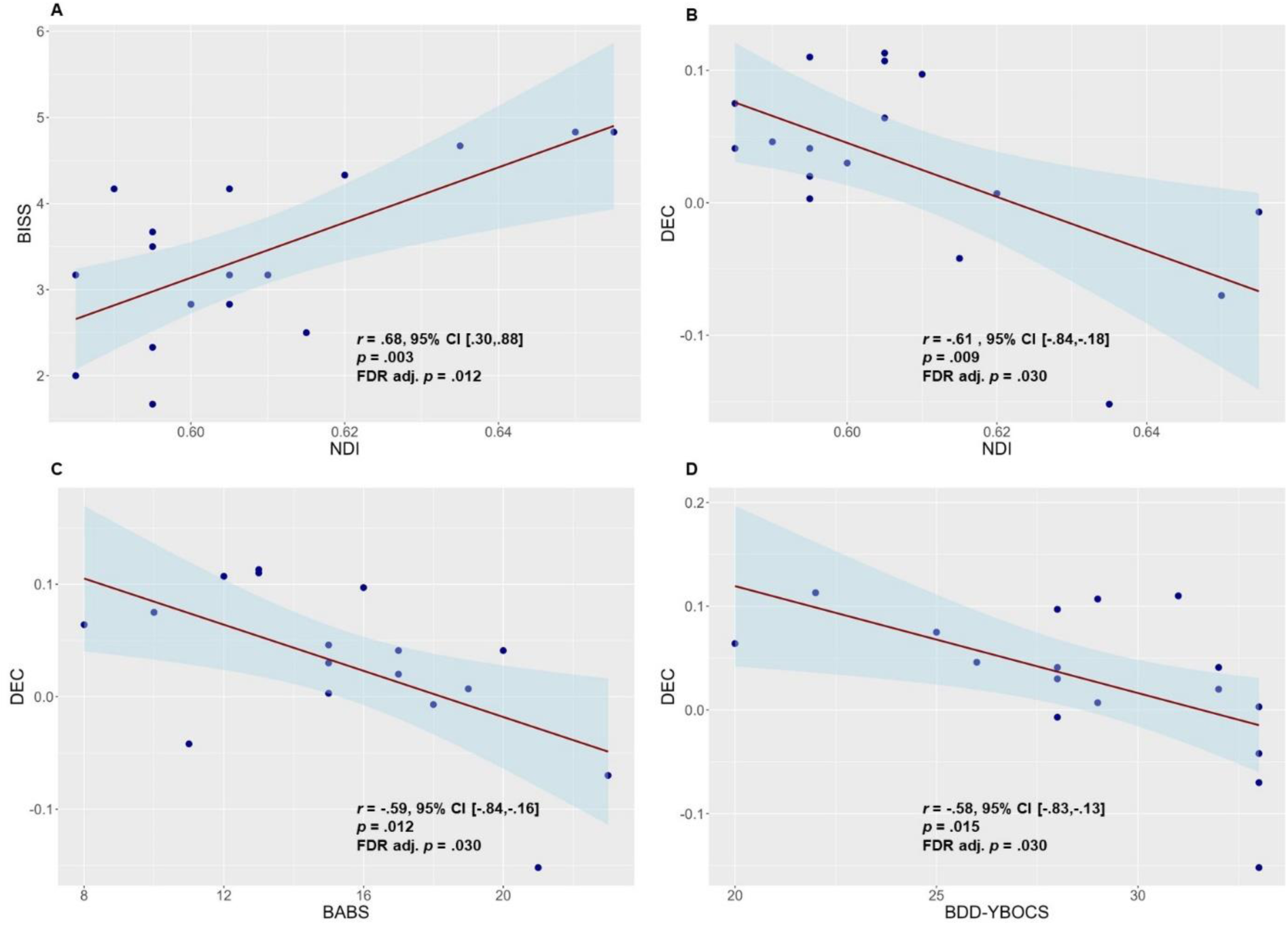
Scatterplots of the significant correlation results. Panels A depicts the significant relationship detected between NDI and BISS from hypothesis testing. Panels B-D depict the significant relationships detected from exploratory analyses including the relationship between NDI and DEC (B), BABS and DEC (C), and BDD-YBOCS and DEC (D).

### 3.4 Post hoc moderation analyses

As *post hoc* analyses to follow up the significant correlations in BDD between NDI and BISS and NDI and DEC, we conducted a moderation analysis of NDI as the predictor of BISS with DEC as a moderator. The model that included NDI as a predictor of BISS with DEC as a moderator was statistically significant (*R*^2^ = 0.52, *F* _(3, 13)_ = 4.62, *p* = .021). This model improved the fit compared to a model without predictors. However, the interaction between NDI and DEC was non-significant (*b* = −237.32, *t* = −1.11, *p* = .287). This suggests that the impact of NDI on BISS remains relatively consistent across different levels of DEC.

## 4 Discussion

The goal of this study was to examine WM microstructure elements in the VOF, the major tract connecting the dorsal and ventral visual streams, and examine their relationships with clinical symptoms that may be associated with visual perceptual distortions in BDD. We additionally explored associations with functional (effective) connectivity from the dorsal to ventral visual systems when viewing one’s face. The main findings were that those with BDD with worse appearance dissatisfaction have a lower fraction of VOF white matter having axons or dendrites. Further, those with a lower fraction of tissue having axons or dendrites had higher connectivity from dorsal to ventral visual areas during face viewing. These functional-structural-appearance evaluation relationships were observed in BDD but not HC. Finally, those with weaker connectivity from dorsal to ventral visual systems had worse BDD symptoms and worse insight. These findings shed light on the inter-relationships between structural and functional connectivity of visual pathway connecting the visual systems responsible for global and local processing in BDD, and their relationships with appearance evaluations and symptomatology.

From our results, worse appearance dissatisfaction in those with BDD was related to lower neurite density along the VOF. Only recently has the importance of VOF integrity in visual processing, and its role in visual processing abnormalities been explored. The VOF has been found in healthy individuals to play a role in stereoscopic depth discrimination, a key aspect of 3D depth perception ^[54, 55]^. Only a few studies have examined the significance of VOF integrity in psychiatric or neurological conditions, and none have previously investigated this in BDD. One study explored VOF WM integrity in multiple sclerosis, a condition known for visual processing abnormalities, finding lower bilateral VOF FA compared to HC ^[56]^. Additionally, a study of individuals with bulimia nervosa, who may also experience perceptual distortion of their appearance ^[57, 58]^ and demonstrate brain activation differences from HC in visual systems when viewing bodies^[59]^, found increased VOF MD^[60]^. Neurite density and orientation dispersion, however, have not yet been examined in the VOF in clinical populations (although, in general, lower neurite density can result in lower FA in some WM tracts^[61]^). In short, relationships between VOF microstructure and general visual functioning and clinical symptomatology have only sparsely been examined.

In general, WM structure influences the transfer of neural electrical signals ^[62]^. Further, better WM integrity, in terms of fractional anisotropy, axial diffusivity, radial diffusivity, and mean diffusivity, is linked to improved performance efficiency, including faster reaction times, across various tasks ^[63–67]^. Thus, it is possible that in the current study those with more disrupted WM integrity could have impaired communication between dorsal and ventral visuals streams, hindering integration of global and local information. This, in addition to the aforementioned reduced DVS activity and connectivity and enhanced VSS activity found in previous studies, might also contribute to imbalances in global and local processing in BDD. This in turn, could result in distorted perception, in turn leading to worse appearance evaluations. In addition, weaker functional connections from the dorsal to the ventral visual stream could also result in poorer integration of global and local information. This could contribute to distorted perceptions that could cause appearance preoccupations and repetitive behaviours (reflected in higher BDD-YBOCS scores). This could also lead to poorer insight (higher BABS scores), as individuals may have difficulty refuting what they are perceiving since most lack knowledge that their visual perception is distorted ^[22]^.

In addition, post hoc exploratory analyses showed that neither anxiety nor depression were significantly associated with effective connectivity or WM microstructure metrics (see Supplemental Information). This adds specificity to the clinical association findings, supporting the notion that appearance dissatisfaction, BDD symptom severity, and insight may be (at least partially) a function of distorted perception.

Relationships were also evident in individuals with BDD between effective connectivity and VOF microstructure metrics. Specifically, higher effective connectivity from dorsal to ventral visual systems during face viewing was associated with lower neurite density. White matter microstructure or diffusion measures have seldom been examined previously in relationship to task-based effective connectivity. One study reported a positive relationship between effectivity connectivity and mean diffusivity between the pre-supplementary motor area and the subthalamic nucleus during an inhibitory control task ^[68]^. Another study found that higher FA of the superior longitudinal fasciculus and cingulum bundle in young adults was associated with greater frontoposterior effective connectivity on an arithmetic interference effect task ^[69]^. Further, in older adults in that study, stronger task-related effective connectivity was linked to higher frontoparietal FA along the inferior fronto-occipital fasciculus. Concerning NDI, a recent systematic review of NODDI measures in psychiatric disorders ^[70]^ did not find any relationships between NDI and effective connectivity, with the authors noting that the extant body of literature on NODDI remains small at this point. In sum, little is thus far known about relationships between diffusion measures and effective connectivity between corresponding regions during tasks, and less about how this might relate to psychopathology.

In the current study, these function/structure relationships were evident in BDD but not HC. Moreover, neither NDI nor effective connectivity were significantly lower in BDD than HC. One speculative explanation for the combination of these findings is that increases in effective connectivity might be compensatory in those with BDD with low neurite density, such that those who are able to increase their effective connectivity (which could result in better global/local integration) have lower BDD symptoms. If so, HC, on the other hand, would not need to engage in this compensation, perhaps explaining why their neurite density and effective connectivity are not significantly related.

There are several limitations of this study to consider. Firstly, as a first-of-its kind study, it has a modest sample size. This affected the power to determine, for example, if significant differences exist between BDD and HC. Thus, a larger study will be important to replicate and confirm the results. In addition, the study population underrepresents the proportion of males with BDD in the general population ^[71, 72]^; thus, findings may not generalize. Another limitation is that the participants’ emotional states during face viewing were not assessed (in the interest of not interrupting natural face viewing processes) such that we could not investigate how degree of emotional arousal, such as state anxiety ^[73]^, affects visual system activity. Since this is a cross-sectional study, a further limitation is that cause and effect relationships cannot be established.

If replication in a larger sample is achieved, this could inform future pre-translational and translational studies. These include, for example, examinations of how targeted interventions, such as attentional or perceptual retraining ^[47]^ affect communication in this tract, and if and how this might change appearance perception. Further, it can inform potential application of non-invasive neuromodulation to the dorsal visual stream (as we have previously piloted with transcranial magnetic stimulation ^[74]^), including providing an early rationale for exploring personalized target site selection ^[75]^ based on white matter tracts or effective connectivity patterns.

## Conclusions

This is the first study to examine the WM microstructure of the VOF in BDD. The results suggest a relationship between disrupted integrity of this tract and worse appearance evaluations, which might be due to impaired communication between visual regions involved in global and local processing. Further, those with weaker effective connectivity between dorsal and ventral visual systems may also have reduced integration of global and local information, possibly contributing to worse perceptual distortions that in turn lead to worse BDD symptoms and worse insight. The results also suggest the presence of distinct functional-structural and perceptual-structural relationships in BDD that are not apparent in HC. In sum, findings in this study provide early insights into how structural integrity of the VOF relates to perceptual distortion-related clinical symptoms in BDD, which may have implications for neurobiologically-based treatment development.

## Acknowledgments

The authors declare no conflicts of interest.

This study was supported by the National Institute of Mental Health (R21MH110865 to JDF, R01MH121520 to JDF), and the Nathan Cumming Foundation (JDF).

## Author Contributions

WW, HP, and JD were responsible for data analysis and paper writing. RC was responsible for modeling white matter microstructure. JDF was responsible for clinical assessment, experimental design, and paper writing. All authors read and approved the submitted manuscript.

## Data Availability

The data reported in this study are available from the corresponding author on request.

## Notes

### Competing Interest Statement

The authors have declared no competing interest.

## References

1. Phillips, K.A., The broken mirror: Understanding and treating body dysmorphic disorder, Rev. & exp ed. The broken mirror: Understanding and treating body dysmorphic disorder, Rev. & exp ed. 2005, New York, NY, US: Oxford University Press. 412, xii, 412-xii.

2. Phillips, K.A. and W. Menard, Suicidality in Body Dysmorphic Disorder: A Prospective Study. American Journal of Psychiatry, 2006. 163(7): p. 1280–1282.

3. Phillips, K.A., et al., A comparison of delusional and nondelusional body dysmorphic disorder in 100 cases. Psychopharmacology bulletin, 1994. 30 **2**: p. 179–86.

4. Veale, D., et al., Body dysmorphic disorder in different settings: A systematic review and estimated weighted prevalence. Body Image, 2016. 18: p. 168–186.

5. Grace, S.A., et al., The neurobiology of body dysmorphic disorder: A systematic review and theoretical model. Neuroscience & Biobehavioral Reviews, 2017. 83: p. 83–96.

6. Li, W., D. Arienzo, and J.D. Feusner, *Body Dysmorphic Disorder: Neurobiological Features and an Updated Model.* Zeitschrift fur klinische Psychologie und Psychotherapie (Gottingen, Germany), 2013. 42(3): p. 184–184.

7. Beilharz, F. and S.L. Rossell, Treatment Modifications and Suggestions to Address Visual Abnormalities in Body Dysmorphic Disorder. Journal of Cognitive Psychotherapy, 2017. 31(4): p. 272–284.

8. Feusner, J.D., et al., Abnormalities of Visual Processing and Frontostriatal Systems in Body Dysmorphic Disorder. Archives of General Psychiatry, 2010. 67(2): p. 197–205.

9. Feusner, J.D., et al., Visual Information Processing of Faces in Body Dysmorphic Disorder. Archives of General Psychiatry, 2007. 64(12): p. 1417–1425.

10. Feusner, J.D., et al., Abnormalities of object visual processing in body dysmorphic disorder. Psychological Medicine, 2011. 41(11): p. 2385–2397.

11. Nassi, J.J. and E.M. Callaway, Parallel processing strategies of the primate visual system. Nature Reviews Neuroscience, 2009. 10(5): p. 360–372.

12. Dhir, S., et al., Parameters of visual processing abnormalities in adults with body image concerns. PLOS ONE, 2018. 13(11): p. e0207585–e0207585.

13. Mundy, M.E. and A. Sadusky, Abnormalities in visual processing amongst students with body image concerns. Advances in Cognitive Psychology, 2014. 10(2): p. 39–39.

14. Feusner, J.D., et al., Inverted face processing in body dysmorphic disorder. Journal of Psychiatric Research, 2010. 44(15): p. 1088–1094.

15. Jefferies, K., K.R. Laws, and N.A. Fineberg, Superior face recognition in Body Dysmorphic Disorder. Journal of Obsessive-Compulsive and Related Disorders, 2012. 1(3): p. 175–179.

16. Stangier, U., et al., Discrimination of Facial Appearance Stimuli in Body Dysmorphic Disorder. Journal of Abnormal Psychology, 2008. 117(2): p. 435–443.

17. Toh, W.L., D.J. Castle, and S.L. Rossell, Face and Object Perception in Body Dysmorphic Disorder versus Obsessive-Compulsive Disorder: The Mooney Faces Task. J Int Neuropsychol Soc, 2017. 23(6): p. 471–480.

18. Li, W., et al., Anorexia nervosa and body dysmorphic disorder are associated with abnormalities in processing visual information. Psychological Medicine, 2015. 45(10): p. 2111–2122.

19. Li, W., et al., Aberrant early visual neural activity and brain-behavior relationships in anorexia nervosa and body dysmorphic disorder. Frontiers in Human Neuroscience, 2015. 9(JUNE): p. 130679–130679.

20. Moody, T.D., et al., Functional connectivity for face processing in individuals with body dysmorphic disorder and anorexia nervosa. Psychological Medicine, 2015. 45(16): p. 3491–3503.

21. Moody, T.D., et al., Brain activation and connectivity in anorexia nervosa and body dysmorphic disorder when viewing bodies: relationships to clinical symptoms and perception of appearance. Brain Imaging and Behavior, 2021. 15(3): p. 1235–1252.

22. Diaz-Fong, J.P. and J.D. Feusner, Visual Perceptual Processing Abnormalities in Body Dysmorphic Disorder. Springer Berlin Heidelberg: Berlin, Heidelberg. p. 1–23.

23. Takemura, H., et al., *A Major Human White Matter Pathway Between Dorsal and Ventral Visual Cortex.* Cerebral Cortex (New York, NY), 2016. 26(5): p. 2205–2205.

24. Yeatman, J.D., et al., The vertical occipital fasciculus: A century of controversy resolved by in vivo measurements. Proceedings of the National Academy of Sciences of the United States of America, 2014. 111(48): p. E5214–E5214.

25. Galletti, C. and P. Fattori, The dorsal visual stream revisited: Stable circuits or dynamic pathways? Cortex, 2018. 98: p. 203–217.

26. Morris, A.P., et al., Dynamics of eye-position signals in the dorsal visual system. Curr Biol, 2012. 22(3): p. 173–9.

27. Siegel, M., et al., Neuronal synchronization along the dorsal visual pathway reflects the focus of spatial attention. Neuron, 2008. 60(4): p. 709–19.

28. Issa, E.B., C.F. Cadieu, and J.J. DiCarlo, Neural dynamics at successive stages of the ventral visual stream are consistent with hierarchical error signals. eLife, 2018. 7: p. e42870.

29. Kravitz, D.J., et al., The ventral visual pathway: an expanded neural framework for the processing of object quality. Trends Cogn Sci, 2013. 17(1): p. 26–49.

30. Takahashi, E., K. Ohki, and D.-S. Kim, Dissociation and convergence of the dorsal and ventral visual working memory streams in the human prefrontal cortex. NeuroImage, 2013. 65: p. 488–498.

31. Tanaka, J.W. and M.J. Farah, Parts and Wholes in Face Recognition. The Quarterly Journal of Experimental Psychology Section A, 1993. 46(2): p. 225–245.

32. Ungerleider, L.G. and J.V. Haxby, ‘What’ and ‘where’ in the human brain. Current Opinion in Neurobiology, 1994. 4(2): p. 157–165.

33. Kircher, T.T.J., et al., Recognizing one’s own face. Cognition, 2001. 78(1): p. B1–B15.

34. Kanwisher, N., J. McDermott, and M.M. Chun, The Fusiform Face Area: A Module in Human Extrastriate Cortex Specialized for Face Perception, in Foundations in Social Neuroscience. 2002, The MIT Press. p. 0.

35. Zachariou, V., et al., Spatial Mechanisms within the Dorsal Visual Pathway Contribute to the Configural Processing of Faces. Cereb Cortex, 2017. 27(8): p. 4124–4138.

36. Zhang, H., et al., NODDI: Practical in vivo neurite orientation dispersion and density imaging of the human brain. NeuroImage, 2012. 61(4): p. 1000–1016.

37. Eisen, J.L., et al., Insight in obsessive compulsive disorder and body dysmorphic disorder. Comprehensive Psychiatry, 2004. 45(1): p. 10–15.

38. Rief, W., et al., The prevalence of body dysmorphic disorder: a population-based survey. Psychological Medicine, 2006. 36(6): p. 877–885.

39. Phillips, K.A., et al., A severity rating scale for body dysmorphic disorder: development, reliability, and validity of a modified version of the Yale-Brown Obsessive Compulsive Scale. Psychopharmacology bulletin, 1997. 33 **1**: p. 17–22.

40. Cash, T.F., et al., Beyond body image as a trait: The development and validation of the body image states scale. Eating Disorders, 2002. 10(2): p. 103–113.

41. Montgomery, S.A. and M. Asberg, A New Depression Scale Designed to be Sensitive to Change. The British Journal of Psychiatry, 1979. 134(4): p. 382–389.

42. Hamilton, M.A.X., THE ASSESSMENT OF ANXIETY STATES BY RATING. British Journal of Medical Psychology, 1959. 32(1): p. 50–55.

43. Jenkinson, M., et al., FSL. NeuroImage, 2012. 62(2): p. 782–790.

44. Cabeen, R.P., D.H. Laidlaw, and A.W. Toga, Quantitative imaging toolkit: software for interactive 3D visualization, data exploration, and computational analysis of neuroimaging datasets. ISMRM-ESMRMB Abstracts, 2018: p. 12–14.

45. Esteban, O., et al., FMRIPrep: a robust preprocessing pipeline for functional MRI. Nature methods, 2019. 16(1): p. 111–111.

46. Wong, W.W., et al., Effects of visual attention modulation on dynamic effective connectivity and visual fixation during own-face viewing in body dysmorphic disorder, in medRxiv. 2021.

47. Wong, W.W., et al., Neural and behavioral effects of modification of visual attention in body dysmorphic disorder. Translational Psychiatry, 2022. 12(1).

48. Wu, G.R., et al., A blind deconvolution approach to recover effective connectivity brain networks from resting state fMRI data. Medical Image Analysis, 2013. 17(3): p. 365–374.

49. Handwerker, D.A., J.M. Ollinger, and M. D’Esposito, Variation of BOLD hemodynamic responses across subjects and brain regions and their effects on statistical analyses. NeuroImage, 2004. 21(4): p. 1639–1651.

50. David, O., et al., Identifying Neural Drivers with Functional MRI: An Electrophysiological Validation. PLOS Biology, 2008. 6(12): p. e315–e315.

51. Büchel, C. and K.J. Friston, Dynamic changes in effective connectivity characterized by variable parameter regression and kalman filtering. Human Brain Mapping, 1998. 6(5-6): p. 403–403.

52. Xia, M., J. Wang, and Y. He, BrainNet Viewer: A Network Visualization Tool for Human Brain Connectomics. PLOS ONE, 2013. 8(7): p. e68910.

53. Igartua, J.J. and A.F. Hayes, Mediation, Moderation, and Conditional Process Analysis: Concepts, Computations, and Some Common Confusions. The Spanish Journal of Psychology, 2021. 24(6): p. e49–e49.

54. Oishi, H., et al., Microstructural properties of the vertical occipital fasciculus explain the variability in human stereoacuity. Proc Natl Acad Sci U S A, 2018. 115(48): p. 12289–12294.

55. Sihvonen, A.J., et al., Structural white matter connectometry of reading and dyslexia. NeuroImage, 2021. 241: p. 118411.

56. Abdolalizadeh, A., S. Mohammadi, and M.H. Aarabi, The forgotten tract of vision in multiple sclerosis: vertical occipital fasciculus, its fiber properties, and visuospatial memory. Brain Structure and Function, 2022. 227(4): p. 1479–1490.

57. Mölbert, S.C., et al., Depictive and metric body size estimation in anorexia nervosa and bulimia nervosa: A systematic review and meta-analysis. Clinical Psychology Review, 2017. 57: p. 21–31.

58. Sattler, F.A., S. Eickmeyer, and J. Eisenkolb, *Body image disturbance in children and adolescents with anorexia nervosa and bulimia nervosa: a systematic review.* Eating and Weight Disorders - Studies on Anorexia, Bulimia and Obesity, 2020. 25(4): p. 857–865.

59. Mohr, H.M., et al., Body image distortions in bulimia nervosa: Investigating body size overestimation and body size satisfaction by fMRI. NeuroImage, 2011. 56(3): p. 1822–1831.

60. Chen, Q., et al., Characteristics of white matter alterations along fibres in patients with bulimia nervosa: A combined voxelwise and tractography study. European Journal of Neuroscience, 2023. 58(3): p. 2874–2887.

61. Beaulieu, C., The Biological Basis of Diffusion Anisotropy. Diffusion MRI, 2009: p. 105–126.

62. Babaeeghazvini, P., et al., Brain Structural and Functional Connectivity: A Review of Combined Works of Diffusion Magnetic Resonance Imaging and Electro-Encephalography. Frontiers in Human Neuroscience, 2021. 15.

63. Bells, S., et al., Changes in White Matter Microstructure Impact Cognition by Disrupting the Ability of Neural Assemblies to Synchronize. J Neurosci, 2017. 37(34): p. 8227–8238.

64. Chung, S., et al., Working Memory And Brain Tissue Microstructure: White Matter Tract Integrity Based On Multi-Shell Diffusion MRI. Scientific Reports, 2018. 8(1): p. 3175.

65. McDonough, I.M. and J.T. Siegel, The Relation Between White Matter Microstructure and Network Complexity: Implications for Processing Efficiency. Frontiers in Integrative Neuroscience, 2018. 12.

66. Rasooli, A., et al., White matter and neurochemical mechanisms underlying age-related differences in motor processing speed. iScience, 2023. 26(6): p. 106794.

67. Roalf, D.R., et al., White matter microstructure in schizophrenia: Associations to neurocognition and clinical symptomatology. Schizophrenia Research, 2015. 161(1): p. 42–49.

68. Rae, C.L., et al., The Prefrontal Cortex Achieves Inhibitory Control by Facilitating Subcortical Motor Pathway Connectivity. The Journal of Neuroscience, 2015. 35(2): p. 786.

69. Hinault, T., et al., Age-related differences in the structural and effective connectivity of cognitive control: a combined fMRI and DTI study of mental arithmetic. Neurobiology of Aging, 2019. 82: p. 30–39.

70. Kraguljac, N.V., et al., Neurite Orientation Dispersion and Density Imaging in Psychiatric Disorders: A Systematic Literature Review and a Technical Note. Biological Psychiatry Global Open Science, 2023. 3(1): p. 10–21.

71. Abujaoude, E., et al., The Prevalence of Body Dysmorphic Disorder in the United States Adult Population. CNS Spectrums, 2008. 13(4): p. 316–322.

72. Taqui, A.M., et al., Body Dysmorphic Disorder: Gender differences and prevalence in a Pakistani medical student population. BMC Psychiatry, 2008. 8: p. 20–20.

73. Bohon, C., et al., Nonlinear relationships between anxiety and visual processing of own and others’ faces in body dysmorphic disorder. Psychiatry research, 2012. 204(2-3): p. 132–132.

74. Wong, W.W., et al., Can excitatory neuromodulation change distorted perception of one’s appearance? Brain Stimulation, 2021. 14(5).

75. Cash, R.F.H. and A. Zalesky, Personalized and Circuit-Based Transcranial Magnetic Stimulation: Evidence, Controversies, and Opportunities. Biological Psychiatry, 2024. 95(6): p. 510–522.

