## Supplementary Materials for "Visual system structural and functional connections during face viewing in body dysmorphic disorder"

**Title**

### Supplementary Methods

1. Exclusion criteria
2. Clinical assessments
3. Image acquisition
4. Anatomical data preprocessing
5. Functional data preprocessing
6. Diffusion data processing
7. Brain connectivity analysis
8. Assumption testing for statistical analyses
9. Post hoc exploratory correlations

### Supplementary Tables

**Table S1** Shapiro-Wilk test of normality for clinical scores in BDD
**Table S2** Shapiro-Wilk test of normality for dynamic effective connectivity dorsal to ventral stream
**Table S3** Shapiro-Wilk test of normality for white matter microstructure of vertical occipital fasciculus

### Supplementary References

### 1. Exclusion criteria

Healthy controls were excluded if they had any current DSM-5 disorder or lifetime bipolar disorder or psychosis. Other exclusion criteria that applied to all participants included psychiatric medications, suicidality, self-injurious behavior, lifetime neurological disorder, current pregnancy, any medical illness that could affect cerebral metabolism, or current treatment with cognitive-behavioral therapy.

### 2. Clinical assessments

***Mini International Neuropsychiatric Interview (MINI):*** The MINI 7.0.0 is a short, structured diagnostic interview developed by psychiatrists and clinicians for DSM-5 and ICD-10 psychiatric disorders. With an administration time of ~15 minutes, the MINI is the structured psychiatric interview of choice for psychiatric evaluation and outcome tracking in clinical psychopharmacology trials and epidemiological studies. The MINI is the most widely used psychiatric structured diagnostic interview instrument in the world, employed by mental health professionals and health organizations in more than 100 countries.

***BDD Diagnostic Module (BDD-DM):*** The BDD-DM is a brief semi-structured diagnostic interview for BDD.

***Yale-Brown Obsessive-Compulsive Scale Modified for BDD (BDD-YBOCS):*** The BDD-YBOCS is a 12-item semi-structured, clinician-rated measure of current BDD severity used in many BDD studies. The first 5 items assess obsessional preoccupations about perceived appearance defects (time preoccupied, interference in functioning and distress due to perceived appearance defects, resistance against preoccupations, and control over preoccupations). Items 6-10 assess BDD-related repetitive behaviors and are similar to items 1-5 (time spent performing the behaviors, interference in functioning due to the behaviors, distress experienced if the behaviors are prevented, and resistance of and control over the behaviors). Item 11 assesses insight into appearance beliefs, and item 12 assesses avoidance because of BDD symptoms. Scores for each item range from 0 (no symptoms) to 4 (extreme symptoms); the total score ranges from 0 to 48, with higher scores reflecting more severe symptoms.

***Brown Assessment of Beliefs Scale (BABS):*** The BABS is a 7-item semi-structured rater-administered scale that assesses insight/delusionality both dimensionally and categorically. BABS items assess the person's conviction that their belief is accurate, perception of others' views of the belief, explanation for any difference between the person's and others' views of the belief, whether the person could be convinced that the belief is wrong, attempts to disprove the belief, insight, and ideas/delusions of reference related to the belief. The first 6 items are summed to create a total score that ranges from 0 to 24; higher scores indicate poorer insight. Item 7 is not included in the total score, because referential thinking is characteristic of some but not all disorders.

***Body Image States Scale (BISS):*** The BISS is a 6-item questionnaire to rate the current body experience: 1) dissatisfaction-satisfaction with one's overall physical appearance, 2) dissatisfaction-satisfaction with one's body size and shape, 3) dissatisfaction-satisfaction with one's weight, 4) feelings of physical attractiveness-unattractiveness, 5) current feelings about one's looks relative to how one usually feels, and 6) evaluation of one's appearance relative to how the average person looks. Responses to each item are based on 9-point, bipolar, Likert-type scales, semantically anchored at each point. The participants were instructed to respond how they felt at that very moment. Higher BISS scores indicate more favorable body image states.

***Montgomery-Åsberg Depression Rating Scale (MADRS):*** The MADRS is a 10-item diagnostic questionnaire which psychiatrists used to measure the severity of depressive episodes. The questionnaire includes questions on the following symptoms: 1) apparent sadness, 2) reported sadness, 3) inner tension, 4) reduced sleep, 5) reduced appetite, 6) concentration difficulties, 7) lassitude, 8) inability to feel, 9) pessimistic thoughts, and 10) suicidal thoughts. Higher score indicates more severe depression, and each item yields a score of 0 to 6. The overall score ranges from 0 to 60.

***Hamilton Anxiety Scale (HAMA):*** The HAMA is a 14-item psychological questionnaire used by clinicians to rate the severity of a patient's anxiety. Each of the 14 items is defined by a series of symptoms, and measures both psychic anxiety (mental agitation and psychological distress) and somatic anxiety (physical complaints related to anxiety). Each item is rated on a scale of 0 (not present) to 4 (severe), with a total score range of 0-56.

### 3. Image acquisition

A 3T Siemens Prisma scanner was used to obtain the MR images. A high-resolution T1-weighted structural image was acquired using ultrafast gradient echo sequence for anatomical reference (TR/TE: 2300/2.27 ms; flip angle: 8°; 256 x 256 matrix; voxel size: 1 x 0.977 x 0.977 mm; 192 slices). Diffusion-weighted images were acquired with 99 directions and three shells (b=0, b=1500, b=3000 s/mm$^2$)(TR/TE: 3222/89.20 ms; flip angle: 78°; voxel size: 1.5 mm$^3$. 140 x 140 matrix; 0 mm gap; 92 slices). Functional images were acquired using a T2*-weighted echo planar imaging (EPI) gradient-echo pulse sequence (TR/TE: 2500/25 ms; flip angle: 80°; 64 x 64 matrix; voxel size: 3 mm$^3$, with a 0.75 mm gap; 34 slices; 124 volumes per run).

### 4. Anatomical data preprocessing

The T1-weighted (T1w) image was corrected for intensity non-uniformity (INU)[1]with ANTs 2.2.0[2], and used as T1w-reference. The T1w-reference was skull-stripped, and brain tissue segmentation of cerebrospinal fluid (CSF), white-matter (WM) and gray-matter (GM) was performed on the brain-extracted T1w using FSL 5.0.9[3]. Brain surfaces were reconstructed using FreeSurfer 6.0.1[4]. Volume-based spatial normalization to standard space was performed through nonlinear registration with ANTs 2.2.0, using brain-extracted versions of both T1w reference and the T1w template (MNI152NLin2009cAsym)[5].

### 5. Functional data preprocessing

For the fMRI run per subject, the following preprocessing was performed. A deformation field to correct for susceptibility distortions was estimated based on fMRIPrep's fieldmap-less approach. The deformation field is that resulting from co-registering the BOLD reference to the same-subject T1w-reference with its intensity inverted[6]. Based on the estimated susceptibility distortion, an unwarped BOLD reference was calculated, and was co-registered to the T1w reference using FreeSurfer 6.0.1 which implements boundary-based registration[7]. Co-registration was configured with nine degrees of freedom to account for distortions remaining in the BOLD reference. Head-motion parameters with respect to the BOLD reference were estimated before any spatiotemporal filtering using FSL 5.0.9[8]. The BOLD timeseries with slice-timing correction were resampled onto their native space by applying a single, composite transform to correct for head-motion and susceptibility distortions. The BOLD timeseries were resampled into standard MNI space, generating the spatially-normalized, preprocessed BOLD runs. Automatic removal of motion artifacts using independent component analysis (ICA-AROMA)[9] was performed on the preprocessed BOLD timeseries on MNI space after removal of non-steady state volumes and spatial smoothing with an isotropic, Gaussian kernel of 6mm FWHM. Corresponding “non-aggressively” denoised runs were produced after such smoothing. Several confounding timeseries were calculated based on the preprocessed BOLD: framewise displacement (FD), DVARS and three region-wise global signals. FD and DVARS were calculated for each run[10]. The three global signals were extracted within the CSF, the WM, and the whole-brain masks. A set of physiological regressors were also extracted to allow for component-based noise correction (CompCor)[11]. Principal components were estimated after high-pass filtering the preprocessed BOLD timeseries for the two CompCor variants: temporal (tCompCor) and anatomical (aCompCor). tCompCor components were calculated from the top 5% variable voxels within a mask covering the subcortical regions. aCompCor components were calculated within the intersection of the aforementioned mask and the union of CSF and WM masks calculated in T1w space. Components were also calculated separately within the WM and CSF masks. For each CompCor decomposition, the k components with the largest singular values were retained, such that the retained components’ timeseries were sufficient to explain 50% of variance across the nuisance mask. The remaining components were dropped from consideration. Frames that exceeded a threshold of 0.5 mm FD or 1.5 standardized DVARS were annotated as motion outliers.

### 6. Diffusion data processing

Diffusion data was preprocessed using FMRIB’s Software Library (FSL v6.0.3; www.fmrib.ox.ac.uk/fsl/) [12]. Echo planar spin-echo (EPI-SE) field maps with opposite phase encoding directions were used to estimate the susceptibility-induced off-resonance field with FSL's TOPUP (as described in [13]). Eddy current-induced distortion correction [14] was applied. Single-subject quality reports were generated using FSL’s EDDY QUAD [15]. Bayesian estimation of diffusion parameters (BEDPOSTX) was used to model and determine the number of crossing fibres per voxel [16].

An atlas-based streamline tractography approach was used to obtain VOF fibre bundle models from the DWI data using the Quantitative Imaging Toolkit (QIT)[17] and other software, noted where applicable. We first manually delineated reference fibre bundles in the IIT ICBM diffusion MRI template [18, 19] using a kernel regression framework [20] for creating a population average diffusion data and using a hybrid probabilistic-deterministic tractography approach [21] for template bundle reconstruction. The reference VOF bundles were segmented according to a priori anatomical information from white matter atlases and reference texts [22, 23], and resulted in seed, inclusion, and exclusion region of interest (ROI) masks for each tract. The diffusion MRI data of each subject was used to compute a deformation from the template to subject space using diffeomorphic registration of the subject FA maps to the IIT template FA maps using Advanced Normalization Tools [2]. The ROI masks for each tract were transformed to subject native space for use in subject-specific bundle reconstruction. We used the tractography module VolumeModelTracksStreamline in QIT to extract bundle models using the bundle masks and the previously estimated ball-and-sticks models. We then reconstructed each VOF bundle using the hybrid probabilistic-deterministic tractography approach [21] with an angle threshold of 75, a minimum volume fraction of 0.05, a step size of 0.5 mm, a minimum length of 10 mm, and trilinear interpolation. For each bundle, seeding was initiated from 5 randomly placed samples within the deformed seed ROI mask, tracks were retained using the deformed inclusion and exclusion ROI masks, and a track density map was computed. We estimated neurite orientation and dispersion imaging (NODDI) parameters from the diffusion MRI data using a spherical mean approach [24]. This resulted in volumetric parameter maps for neurite density index (NDI) and orientation dispersion index (ODI), which were averaged across all bundle vertices to produce summary statistics of VOF microstructure for each hemisphere in each subject. This entire process as automated using the qitdiff command available in QIT.

### 7. Brain connectivity analysis

A similar analysis strategy has been adopted in our previous study for exploring dynamic connectivity for fear processing in BDD and Anorexia Nervoa[25]. We used dynamic effective connectivity (DEC) in this study because of its ability to estimate causal connectivity across time with a precision of each time point, which helped us capture connectivity only within task blocks of interest. DEC was estimated in a Kalman filter based dynamic multivariate vector autoregressive (dMVAR) framework that is based on the concept of Granger causality (GC). Simulations [26, 27] as well as experimental results from electrophysiology and optogenetics [28-31] have demonstrated the ability of GC in estimating fMRI effective connectivity, when latent neural time series data are used after HRF deconvolution (as done in this study).

FMRI is an indirect measure of neural activity corrupted by blood hemodynamics that can alter the relative phase between fMRI time series without having any neural delays between the corresponding regions [32, 33]. Since this phenomenon can confound fMRI connectivity estimates[34], deconvolution was performed. Additionally, deconvolution has also been found to improve GC estimation [28, 30, 35]. A blind-deconvolution technique developed by Wu et al. [36] was used, which is a data-driven technique based on the point process model[37]. It identifies neural events in fMRI data as point processes based on relative intensities and estimates the best fit double-gamma HRF in a least squares sense. Latent neuronal time series is then estimated using Wiener deconvolution. Several studies [38-40] have employed this technique. The deconvolution was performed using rsHRF toolbox implemented in SPM12 (https://github.com/compneuro-da/rsHRF).

DEC was computed at each time point using Kalman-filter based time-varying GC[41]. The basic concept is that if past values of a timeseries can forecast the future values of another timeseries, a causal influence from the former to the latter is inferred. A GC value of 0 represents no causal relationship from source to target timeseries, a value of 1 represents strong positive causality (i.e. increase in BOLD response of the source enhances BOLD response of the target, and vice versa), and a value of -1 represents strong negative causality (i.e. increase in BOLD response of the source suppresses BOLD response of the target, and vice versa)[25]. The deconvolved timeseries were fitted into a dMVAR model for estimating DEC between ROIs, as in prior studies [42, 43]. Further details and underlying mathematics are elaborated in studies [34, 44] . The dMVAR model coefficients vary as a function of time, whose lengths were identical to the number of timepoints in the timeseries. That is, a DEC value was obtained for every timepoint for every connection. Using this, we obtained the desired block-specific connectivity values and aggregated them to represent the corresponding connection[45]. The timepoints associated with those trials of viewing unaltered faces were extracted for subsequent statistical analysis.

***Repetition time issue in dynamic connectivity analysis***

A limitation in this study is the poor repetition time (TR) of 2.5s, which is barely sufficient to capture effective connectivity in the brain[46]. With this modest TR, the data is not sensitive enough to capture causal interactions happening at faster timescales. In other words, a shorter TR would have enabled us to capture more DEC patterns than with a longer TR (or a longer TR could only capture a subset of DEC patterns compared with the patterns captured with a shorter TR). It is possible that there are within-group and/or between-group differences not detected with the existing TR of 2.5s, leading to false negative errors (where the test results incorrectly fail to indicate the presence of a condition when it is present). But importantly, the positive results of this study are still reliable as it does not imply that the positive results captured at a longer TR are less valid compared to what would have been captured at a shorter TR. These aspects must be kept in mind while interpreting negative findings of this study. Future studies must prioritize faster acquisitions in their study designs and tradeoffs so that maximum benefit is drawn from dynamic time series analysis techniques (such as those used in this study).

### 8. Assumption testing for statistical analyses

See tables below for tests of normality. A slight deviation from normality was detected for the NDI distribution in the BDD group. No extreme outliers were detected, skewness (1.19 ± 0.55) and kurtosis (0.65 ± 1.06) were within an acceptable range, and both the independent samples *t*-test and Pearson's correlation are considered robust to minor violations of normality. Given these considerations, and for the purposes of consistency across all tests, we did not transform the variable or perform a non-parametric test.

### 9. Post hoc exploratory correlations

Exploratory correlations between anxiety (HAMA) and depression (MADRS) with WM metrics and DEC were all non-significant: HAMA and NDI ($r$ = .17, $p$ = .317), HAMA and ODI ($r$ = -.16, $p$ = .352), HAMA and DEC ($r$ = -.11, $p$ = .514), MADRS and NDI ($r$ = .15, $p$ = .385), MADRS and ODI ($r$ = -.17, $p$ = .318), MADRS and DEC ($r$ = -.07, $p$ = .657).

**Table S1** Shapiro-Wilk test of normality for clinical scores in BDD

| Group | P-values from Shapiro-Wilk test | | |
|---|---|---|---|
| | BDD-YBOCS | BABS | BISS |
| BDD | .055 | .999 | .558 |

Abbreviations: BDD = body dysmorphic disorder; BDD-YBOCS = Yale-Brown Obsessive-Compulsive Scale Modified for BDD; BABS = Brown Assessment of Beliefs Scale; BISS = Body Image States Scale.

**Table S2** Shapiro-Wilk test of normality for dynamic effective connectivity from dorsal to ventral stream

| Group | P-values from Shapiro-Wilk test |
|---|---|
| BDD | .151 |
| CON | .395 |

Abbreviations: BDD = body dysmorphic disorder; CON = control.

**Table S3** Shapiro-Wilk test of normality for white matter microstructure of vertical occipital fasciculus

| Group | P-values from Shapiro-Wilk test | |
|---|---|---|
| | NDI | ODI |
| BDD | .018 | .062 |
| CON | .089 | .120 |

Abbreviations: BDD = body dysmorphic disorder; CON = control; NDI = neurite density index; ODI = orientation dispersion index; FA = fractional anisotropy; MD = mean diffusivity.

### References


1. Tustison, N.J., et al., *N4ITK: Improved N3 bias correction.* IEEE Transactions on Medical Imaging, 2010. **29**(6): p. 1310-1320.
2. Avants, B.B., et al., *Symmetric diffeomorphic image registration with cross-correlation: Evaluating automated labeling of elderly and neurodegenerative brain.* Medical Image Analysis, 2008. **12**(1): p. 26-41.
3. Zhang, Y., M. Brady, and S. Smith, *Segmentation of brain MR images through a hidden Markov random field model and the expectation-maximization algorithm.* IEEE Transactions on Medical Imaging, 2001. **20**(1): p. 45-57.
4. Dale, A.M., B. Fischl, and M.I. Sereno, *Cortical Surface-Based Analysis: I. Segmentation and Surface Reconstruction.* NeuroImage, 1999. **9**(2): p. 179-194.
5. Fonov, V.S., et al., *Unbiased nonlinear average age-appropriate brain templates from birth to adulthood.* NeuroImage, 2009. **47**: p. S102-S102.
6. Wang, S., et al., *Evaluation of field map and nonlinear registration methods for correction of susceptibility artifacts in diffusion MRI.* Frontiers in Neuroinformatics, 2017. **11**: p. 238590-238590.
7. Greve, D.N. and B. Fischl, *Accurate and robust brain image alignment using boundary-based registration.* NeuroImage, 2009. **48**(1): p. 63-72.
8. Jenkinson, M., et al., *Improved Optimization for the Robust and Accurate Linear Registration and Motion Correction of Brain Images.* NeuroImage, 2002. **17**(2): p. 825-841.
9. Pruim, R.H.R., et al., *ICA-AROMA: A robust ICA-based strategy for removing motion artifacts from fMRI data.* NeuroImage, 2015. **112**: p. 267-277.
10. Power, J.D., et al., *Methods to detect, characterize, and remove motion artifact in resting state fMRI.* NeuroImage, 2014. **84**: p. 320-341.
11. Behzadi, Y., et al., *A component based noise correction method (CompCor) for BOLD and perfusion based fMRI.* NeuroImage, 2007. **37**(1): p. 90-101.
12. Smith, S.M., et al., *Advances in functional and structural MR image analysis and implementation as FSL.* Neuroimage, 2004. **23 Suppl 1**: p. S208-19.
13. Andersson, J.L., S. Skare, and J. Ashburner, *How to correct susceptibility distortions in spin-echo echo-planar images: application to diffusion tensor imaging.* Neuroimage, 2003. **20**(2): p. 870-88.
14. Andersson, J.L.R. and S.N. Sotiropoulos, *An integrated approach to correction for off-resonance effects and subject movement in diffusion MR imaging.* NeuroImage, 2016. **125**: p. 1063-1078.
15. Bastiani, M., et al., *Automated quality control for within and between studies diffusion MRI data using a non-parametric framework for movement and distortion correction.* Neuroimage, 2019. **184**: p. 801-812.
16. Hernández, M., et al., *Accelerating Fibre Orientation Estimation from Diffusion Weighted Magnetic Resonance Imaging Using GPUs.* PLOS ONE, 2013. **8**(4): p. e61892.
17. Cabeen, R.P., D.H. Laidlaw, and A.W. Toga, *Quantitative imaging toolkit: software for interactive 3D visualization, data exploration, and computational analysis of neuroimaging datasets.* ISMRM-ESMRMB Abstracts, 2018: p. 12-14.
18. Varentsova, A., S. Zhang, and K. Arfanakis, *Development of a high angular resolution diffusion imaging human brain template.* Neuroimage, 2014. **91**: p. 177-86.
19. Zhang, S., et al., *Enhanced ICBM diffusion tensor template of the human brain.* Neuroimage, 2011. **54**(2): p. 974-84.
20. Cabeen, R.P., M.E. Bastin, and D.H. Laidlaw, *Kernel regression estimation of fiber orientation mixtures in diffusion MRI.* NeuroImage, 2016. **127**: p. 158-172.
21. Cabeen, R. and A.W. Toga. *Reinforcement Tractography: A Hybrid Approach for Robust Segmentation of Complex Fiber Bundles*. in *2020 IEEE 17th International Symposium on Biomedical Imaging (ISBI)*. 2020.

22. Catani, M., et al., *Virtual in Vivo Interactive Dissection of White Matter Fasciculi in the Human Brain.* NeuroImage, 2002. **17**(1): p. 77-94.
23. Wakana, S., et al., *Reproducibility of quantitative tractography methods applied to cerebral white matter.* NeuroImage, 2007. **36**(3): p. 630-644.
24. Cabeen, R., F. Sepehrband, and A.W. Toga, *Rapid and accurate NODDI parameter estimation with the spherical mean technique.* . Proc International Society for Magnetic Resonance in Medicine (ISMRM) 2019. **3365**.
25. Rangaprakash, D., et al., *Aberrant Dynamic Connectivity for Fear Processing in Anorexia Nervosa and Body Dysmorphic Disorder.* Frontiers in Psychiatry, 2018. **9**: p. 372157-372157.
26. Ryali, S., et al., *Multivariate dynamical systems models for estimating causal interactions in fMRI.* NeuroImage, 2011. **54**(2): p. 807-823.
27. Wen, X., G. Rangarajan, and M. Ding, *Is Granger Causality a Viable Technique for Analyzing fMRI Data?* PLOS ONE, 2013. **8**(7): p. e67428-e67428.
28. David, O., et al., *Identifying Neural Drivers with Functional MRI: An Electrophysiological Validation.* PLOS Biology, 2008. **6**(12): p. e315-e315.
29. Katwal, S.B., et al., *Measuring relative timings of brain activities using fMRI.* NeuroImage, 2013. **66**: p. 436-448.
30. Ryali, S., et al., *Combining optogenetic stimulation and fMRI to validate a multivariate dynamical systems model for estimating causal brain interactions.* NeuroImage, 2016. **132**: p. 398-405.
31. Wang, Y., et al., *Experimental Validation of Dynamic Granger Causality for Inferring Stimulus-Evoked Sub-100 ms Timing Differences from fMRI.* IEEE Transactions on Neural Systems and Rehabilitation Engineering, 2017. **25**(6): p. 539-546.
32. Aguirre, G.K., E. Zarahn, and M. D'Esposito, *The Variability of Human, BOLD Hemodynamic Responses.* NeuroImage, 1998. **8**(4): p. 360-369.
33. Handwerker, D.A., J.M. Ollinger, and M. D'Esposito, *Variation of BOLD hemodynamic responses across subjects and brain regions and their effects on statistical analyses.* NeuroImage, 2004. **21**(4): p. 1639-1651.
34. Rangaprakash, D., et al., *Identifying disease foci from static and dynamic effective connectivity networks: Illustration in soldiers with trauma.* Human Brain Mapping, 2018. **39**(1): p. 264-287.
35. Ryali, S., et al., *Estimation of functional connectivity in fMRI data using stability selection-based sparse partial correlation with elastic net penalty.* NeuroImage, 2012. **59**(4): p. 3852-3861.
36. Wu, G.R., et al., *A blind deconvolution approach to recover effective connectivity brain networks from resting state fMRI data.* Medical Image Analysis, 2013. **17**(3): p. 365-374.
37. Saad, Z.S., et al., *Trouble at Rest: How Correlation Patterns and Group Differences Become Distorted After Global Signal Regression.* https://home.liebertpub.com/brain, 2012. **2**(1): p. 25-32.
38. Amico, E., et al., *Posterior Cingulate Cortex-Related Co-Activation Patterns: A Resting State fMRI Study in Propofol-Induced Loss of Consciousness.* PLOS ONE, 2014. **9**(6): p. e100012-e100012.
39. Boly, M., et al., *Stimulus Set Meaningfulness and Neurophysiological Differentiation: A Functional Magnetic Resonance Imaging Study.* PLOS ONE, 2015. **10**(5): p. e0125337-e0125337.
40. Lamichhane, B., et al., *The Neural Basis of Perceived Unfairness in Economic Exchanges.* https://home.liebertpub.com/brain, 2014. **4**(8): p. 619-630.
41. Büchel, C. and K.J. Friston, *Dynamic changes in effective connectivity characterized by variable parameter regression and kalman filtering.* Human Brain Mapping, 1998. **6**(5-6): p. 403-403.
42. Feng, C., et al., *Diffusion of responsibility attenuates altruistic punishment: A functional magnetic resonance imaging effective connectivity study.* Human Brain Mapping, 2016. **37**(2): p. 663-677.
43. Hutcheson, N.L., et al., *Effective connectivity during episodic memory retrieval in schizophrenia participants before and after antipsychotic medication.* Human Brain Mapping, 2015. **36**(4): p. 1442-1457.
44. Rangaprakash, D., et al., *Dynamics of segregation and integration in directional brain networks: Illustration in soldiers with PTSD and neurotrauma.* Frontiers in Neuroscience, 2019. **13**(JUL): p. 442861-442861.

45. Sathian, K., G. Deshpande, and R. Stilla, *Neural Changes with Tactile Learning Reflect Decision-Level Reweighting of Perceptual Readout.* Journal of Neuroscience, 2013. **33**(12): p. 5387-5398.
46. Abler, B., et al., *Investigating directed influences between activated brain areas in a motor-response task using fMRI*. Magnetic Resonance Imaging, 2006. **24**(2): p. 181-185.
